# Loss of the Krüppel-like factor 4 tumor suppressor is associated with epithelial-mesenchymal transition in colorectal cancer

**DOI:** 10.1101/743443

**Authors:** Kimberley C. Agbo, Jessie Z. Huang, Amr M. Ghaleb, Jennie L. Williams, Kenneth R. Shroyer, Agnieszka B. Bialkowska, Vincent W. Yang

## Abstract

Colorectal cancer (CRC) is the third leading cancer-related cause of death due to its propensity to metastasize. Epithelial-mesenchymal transition (EMT) is a multistep process important for invasion and metastasis of CRC. Krüppel-like factor 4 (KLF4) is a zinc finger transcription factor highly expressed in differentiated cells of the intestinal epithelium. KLF4 has been shown to play a tumor suppressor role during CRC tumorigenesis - its loss accelerates development and progression of cancer. The present study examines the relationship between KLF4 and markers of EMT in CRC.

**Methods:** Immunofluorescence staining for KLF4 and EMT markers was performed on archived patient samples after colorectal cancer resection and on colonic tissues of mice with colitis-associated cancer.

**Results:** We found that KLF4 expression is lost in tumor sections obtained from CRC patients and in those of mouse colon following azoxymethane and dextran sodium sulfate (AOM/DSS) treatment when compared to their respective normal appearing mucosa. Importantly, in CRC patient tumor sections we observed a negative correlation between KLF4 levels and mesenchymal markers including TWIST, β-catenin, claudin-1, N-cadherin, and vimentin. Similarly, in tumor tissues from AOM/DSS-treated mice KLF4 levels were negatively correlated with mesenchymal markers including SNAI2, β-catenin, and vimentin and positively correlated with the epithelial marker E-cadherin.

**Conclusion:** These findings suggest that the loss of KLF4 expression is a potentially significant indicator of EMT in CRC.

## Introduction

Colorectal cancer (CRC) is the third leading cause of the cancer-related deaths often with metastasis to the liver, lung and bone (1, 2). Epithelial-mesenchymal transition (EMT) is a transdifferentiation process that allows a polarized epithelial cell assume a mesenchymal cell phenotype, characterized by enhanced migratory and invasiveness capacity, elevated resistance to apoptosis, and increased synthesis of extracellular matrix (ECM) components (3). During the EMT process, epithelial cells lose apical-basal polarity that is accompanied by reorganization of cytoskeleton and reprogramming of the signaling pathways that allow for an increase in motility and the development of an invasive phenotype. This multistep complex process is characterized by modifications in the expression of a host of transcription factors and specific cell-surface proteins, as well as reorganization and expression of cytoskeletal proteins, and production of enzymes that degrade the ECM (3). A change in some of these factors, such as upregulation of TWIST, SNAI1, SNAI2, ZEB1, vimentin and N-cadherin, and downregulation of E-cadherin and tight junction proteins such as ZO-1 is indicative of progression of EMT (4, 5). In CRC, EMT has been strongly associated with the invasive and metastatic phenotype, thereby generating the life-threatening manifestations of metastatic disease cancer. The activation of EMT program has been suggested to be the critical mechanism for the acquisition of malignant phenotypes by epithelial cancer cells (3).

Krüppel-like factor 4 (KLF4) belongs to the family of zinc-finger transcriptions factors that play critical roles during development, proliferation, differentiation, homeostasis, as well as development and progression of many diseases including inflammation and carcinogenesis (6–8). In the digestive, tract KLF4 is predominantly expressed in differentiated cells of the villus and surface epithelium of the small intestine and colon, respectively (9–13). Importantly, evidence indicates that KLF4 functions as a tumor suppressor during CRC tumorigenesis (12). It has been shown that loss of KLF4 expression is associated with the early stage of CRC development and that KLF4 is a prognostic indicator for CRC survival and recurrence (14, 15). Recently, we demonstrated that KLF4 also plays a protective role against tumor formation during inflammation-induced colorectal tumorigenesis (16–18). The biochemical mechanisms triggering the acquisition of the invasive phenotype and the subsequent systemic spread of the cancer cell have been areas of intensive research. However, the potential role of KLF4 in the regulation of colorectal cancer metastasis has not been addressed. Here, we demonstrate that KLF4’s role in colorectal tumorigenesis extends to its ability to regulate EMT.

## Materials and Methods

### Samples from patients

Surgical specimens of resected colorectal cancer specimen obtained from Stony Brook University and SUNY Downstate were used in this study. A total of 12 specimens were processed for immunofluorescence. All samples were of Caucasian origin, 2 female and 10 male. One sample was qualified as stage I, one – stage 2, two – stage 3, and eight as stage IV. The protocol for the sample collection has been originally approved by the Institutional Review Board by the State University of New York at Stony Brook on October 17^th^, 2014 (CORIHS 2014-2821-F) and qualified for a waiver under the Federal Law of Department of Health and Human Services (DHHS) per article 45CFR46.116.d.

### Mice

All animal studies were approved by the Stony Brook University Institutional Animal Care and Use Committee (IACUC) and performed in accordance with institutional policies and NIH guidelines. Mice with the floxed *Klf4* gene (*Klf4^fl/fl^*) were described previously (12). These mice were derived from a C57BL/6 background and are indistinguishable from the wild-type mice.

### Azoxymethane and dextran sodium sulfate treatment

Azoxymethane (AOM) and dextran sodium sulfate (DSS) treatment has been performed as described previously (18).

### Tissue harvesting and tumor assessment, preparation and immunostaining

Tissues were collected as described previously and were prepared for immunofluorescence as described previously (18). Briefly, tissue sections were baked in a 65°C oven overnight, deparaffinized in xylene, and rehydrated by incubation in a decreasing ethanol bath series (100%, 95%, and 70%), followed by antigen retrieval in citrate buffer solution (10 mM sodium citrate, 0.05% Tween-20, pH 6.0) at 120°C for 10 minutes using a decloaking chamber (Biocare Medical) and 30 minutes incubation at 4°C. The histological sections were incubated with blocking buffer (5% bovine serum albumin and 0.01% Tween 20 in 1X Tris-buffered phosphate-buffered saline (TTBS)) for 1 hour at 37°C. The primary antibodies goat anti-KLF4 (1:200 for human sections and 1:300 for mice sections; R&D: AF3158); mouse anti-PanCK (1:200 for human sections; Biocare Medical: AE1/AE3); rabbit anti-β-catenin (1:500 for human sections and 1:150 for mice sections; Cell Signaling: 8480); rabbit anti-TWIST (1:500; Abcam: ab49254); rabbit anti-Claudin-1 (1:500; Cell Signaling: 13255); rabbit anti-N-cadherin (1:500; Cell Signaling: 13116); rabbit anti-E-cadherin (1:300 for mice sections, Cell Signaling: 3195), rabbit anti-Vimentin (1:500 for human sections and 1:100 for mice sections; Cell Signaling: 5741); and rabbit anti-SNAI2 (1:500 for human sections and 1:400 for mice sections, Cell Signaling: 9585) were added and incubated at 4°C overnight. For KLF4, secondary unconjugated bovine anti-goat antibody was added at 1:500 dilution in blocking buffer for 30 minutes at 37°C. For human sections to stain for PanCK, secondary unconjugated chicken anti-mouse antibody was added at 1:500 dilution in blocking buffer for 30 minutes at 37°C. Appropriate Alexa Fluor–labeled antibodies (Molecular Probes) were added at 1:500 dilution in blocking buffer for 30 minutes at 37°C. For mice sections, a mouse anti-PanCK antibody conjugated with Alexa 488 (1:100; ThermoFisher Scientific: 53-9003-82) were used. All slides were counterstained with Hoechst 33258 (ThermoFisher Scientific: H3569) and mounted with Fluoromount Aqueous Mounting Medium (Sigma-Aldrich: F4680). Slides were analyzed using a Nikon eclipse 90i microscope (Nikon Instruments Inc.) equipped with DS-Qi1Mc and DS-Fi1 CCD cameras (Nikon Instruments Inc.).

### Statistical analysis

Student’s paired or unpaired t test was used for statistical analyses. Differences between values were considered significant when p < 0.05. This analysis was performed using GraphPad Prism version 5.00 for Windows (GraphPad Software, San Diego, CA).

## Results

### Expression of KLF4 in human colorectal cancer is negatively correlated with markers of EMT

Epithelial-mesenchymal transition (EMT) is a precisely orchestrated multistep process regulated by several transcriptional master regulators including TWIST, SNAI1, SNAI2, and ZEB1 (3). To determine the correlation between KLF4 expression and EMT in colorectal cancer (CRC) we performed immunofluorescence analysis of matched pairs of archived samples from patients after tumors’ resections. Firstly, we analyzed the expression pattern of KLF4 and TWIST. As shown in Figure 1 (A, SB396N) KLF4 is expressed in the nucleus of epithelial cells in the normal-appearing mucosa adjacent to the cancer tissues. These cells are positive for PanCK, an epithelial marker, and negative for the nuclear expression of the biomarker of EMT, TWIST. In contrast, in the tumor samples from the same patient (Fig. 1A, SB396T), the expression of KLF4 is downregulated in the epithelial cells, which is accompanied by a significant increase in the expression of TWIST. Our statistical analysis showed that there is a negative correlation between KLF4 and TWIST expression in normal-appearing mucosa and tumor tissues (p < 0.001). Several common signaling pathways regulate factors involved in EMT including HH, WNT, NOTCH and TGF-β (19). WNT signaling plays an important role in the homeostasis of the intestinal epithelium and its deregulation leads to cancer formation which is accompanied by modification of the pattern and level of expression of its major effector, β-catenin (20–22). In the normal-appearing mucosa, β-catenin is predominantly localized to the membrane of the epithelial cells with a modest nuclear staining (Fig. 1B, SB474N). Upon loss of KLF4 in the colorectal tumor (Fig. 1B, SB474T), the levels of the cytoplasmic and nuclear β-catenin are significantly increased while its membranous expression is decreased (p < 0.001). Loss of cell polarity and cell-cell junctions is another hallmark of EMT and is characterized by increased expression of claudin-1 and N-cadherin (23–27). Immunofluorescence staining of claudin-1 (Fig. 2A, SB474N) and N-cadherin (Fig. 2B, SB378N) within the normal-appearing mucosa shows little or no staining while there is pronounced nuclear KLF4 staining. In matching colorectal cancer tumor tissues with lack of KLF4 expression, both claudin-1 and N-cadherin levels are significantly increased (Fig. 2A, SB474T and Fig. 2B, SB378T, respectively). Our analysis showed that there is a significant negative correlation between expression of KLF4 and claudin-1 and N-cadherin between the normal-appearing mucosa and tumor tissues (p < 0.001). Furthermore, staining for vimentin, another mesenchymal marker, showed a lack of expression within the epithelial component of the normal-appearing mucosa (Fig. 3 SB474N), but a slight increase within the epithelial compartment upon loss of KLF4 in the tumor tissues (Fig. 3 SB474T, white arrowheads). Statistical analysis showed that the expression levels of KLF4 and vimentin are negatively correlated in normal mucosa (p < 0.05) and in tumor sections (p < 0.001).

**Figure 1.**
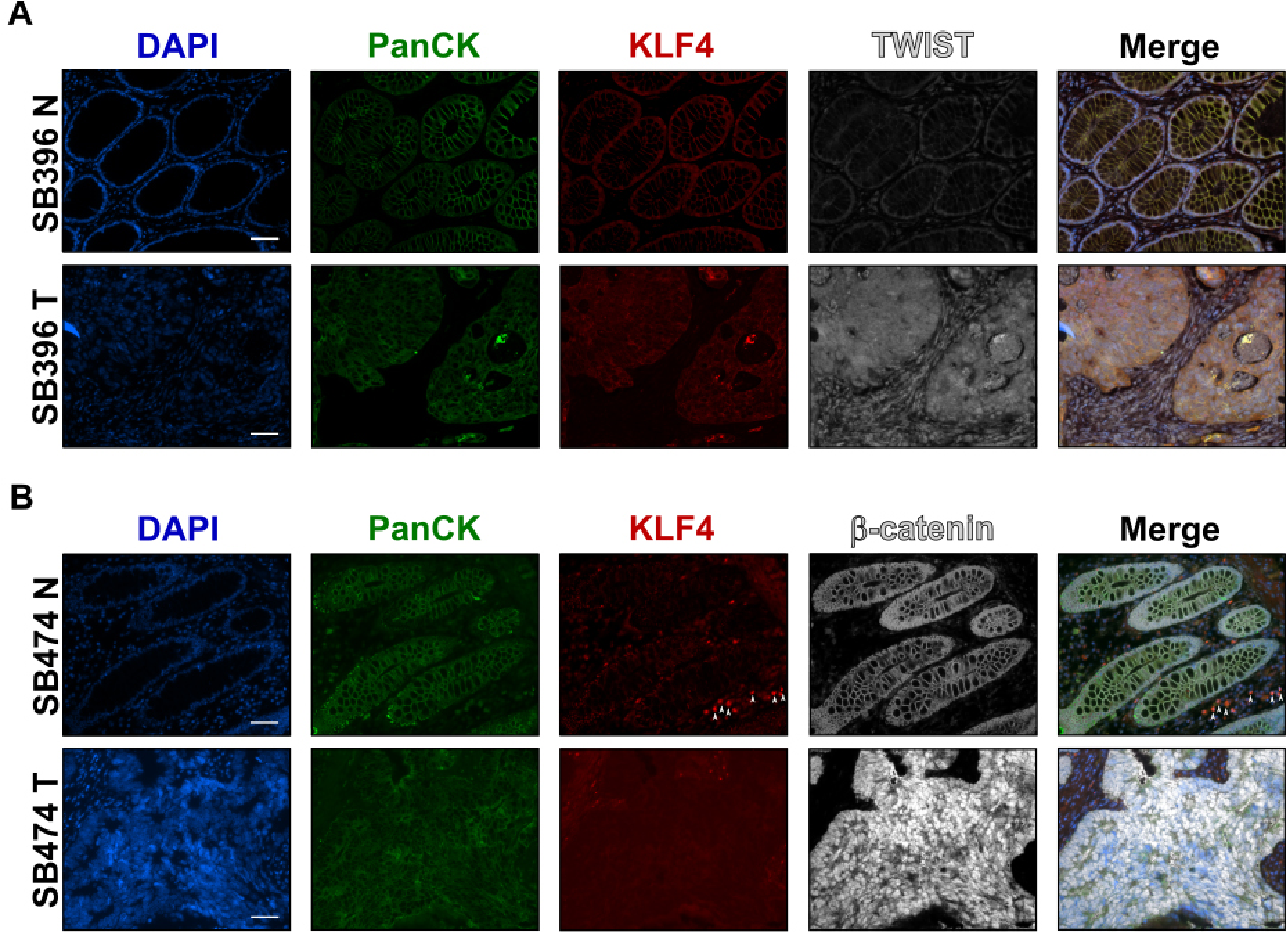
Immunofluorescence staining for PanCK, KLF4, TWIST, and β-catenin in human colonic tissues. Representative images of normal adjacent mucosa and tumor sections from two different human specimens. (**A**) – TWIST (SB396N and SB396T) and (**B**) - β-catenin (SB474N and SB474T). White arrowheads labelled KLF4 stain in stromal tissue. Scale bar, 50 μm.

**Figure 2.**
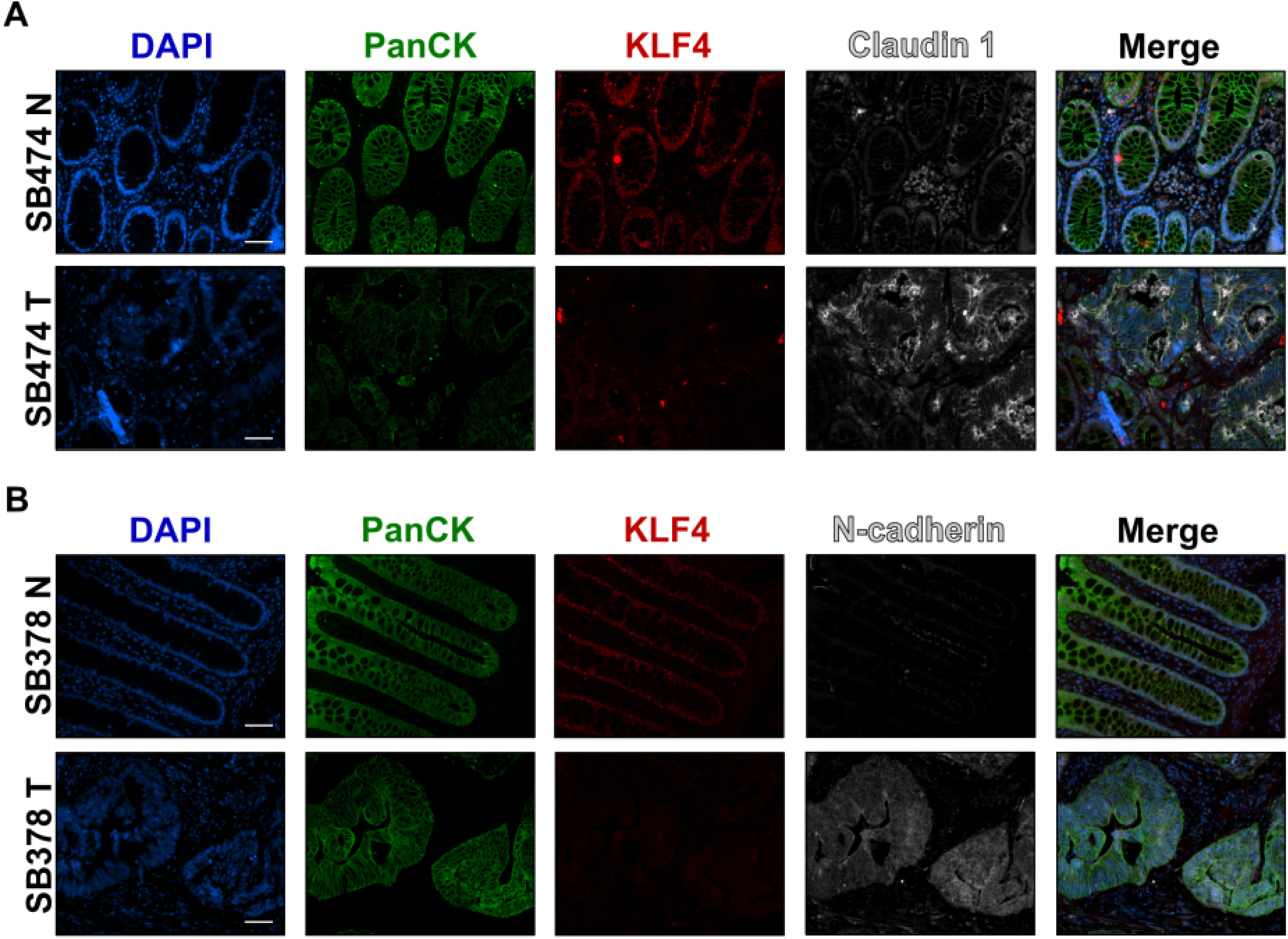
Immunofluorescence staining for PanCK, KLF4, claudin-1, and N-cadherin in human colonic tissues. Representative images of normal adjacent mucosa and tumor sections from two different human specimens. (**A**) – claudin-1 (SB474N and SB474T) and (**B**) – N-cadherin (SB378N and SB378T). Scale bar, 50 μm.

**Figure 3.**
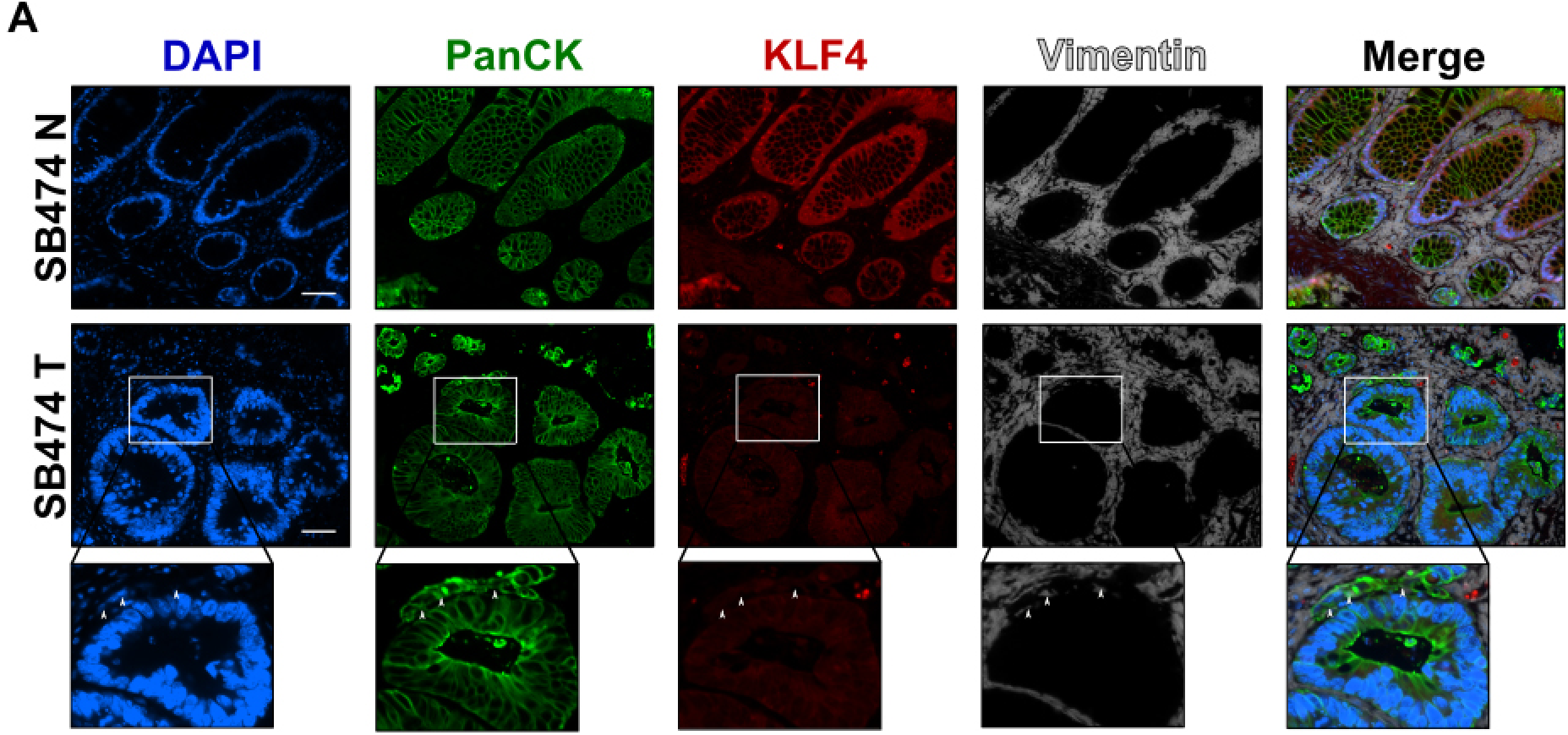
Immunofluorescence staining for PanCK, KLF4, and vimentin in human colonic tissues. Representative images of normal adjacent mucosa (SB474N) and tumor sections (SB474T) from two different human specimens. White arrowheads labelled vimentin stain within epithelial cells. Scale bar, 50 μm.

### KLF4 expression is negatively correlated with markers of EMT in a mouse model of colitis-associated cancer

Recently, in a mouse model, we demonstrated that KLF4 plays a protective role against progression of colitis-associated cancer and that decrease in KLF4 levels is associated with increased aggressiveness of the disease (18). To test if KLF4 expression levels correlate with the expression pattern of EMT markers, we performed immunofluorescence staining of KLF4 and select EMT markers on the mouse tissues (*Klf4^fl/fl^* mice) after treatment with AOM/DSS as described in the Material and Methods section. Immunofluorescence staining showed that in normal mucosa KLF4 is expressed in the nuclei of epithelial cells defined by staining for PanCK. Co-staining for SNAI2, showed that SNAI2 was predominantly expressed in the stromal section of the normal-appearing mucosa and tumor section of mice after AOM/DSS treatment (Fig. 4A) and was absent from epithelial compartment. However, in the tumor section where KLF4 loss was observed, we noticed increased levels of SNAI2 in the nuclei of the epithelial cells that were defined by positive staining for PanCK (Fig. 4A). As in human specimens, we performed immunostaining analysis for β-catenin. However, we did not observe an increase in nuclear β-catenin staining in the tumor sections in comparison to the normal adjacent mucosa (Fig. 4B). With respect to adherens junction complexes, we observed a reduction in the expression levels of E-cadherin in the tumor section as compared to the normal adjacent mucosa (Fig. 5A). Expression of the mesenchymal marker, vimentin in normal adjacent mucosa of mice after AOM/DSS treatment is confined to the stroma (Fig. 5B) and does not correlate with KLF4 expression. However, in the tumor of mice after AOM/DSS treatment, we observed, a slight increase in the staining of vimentin within the epithelial section which suggests a change in the characteristics of these cells from epithelial toward mesenchymal phenotype (Fig. 5B).

**Figure 4.**
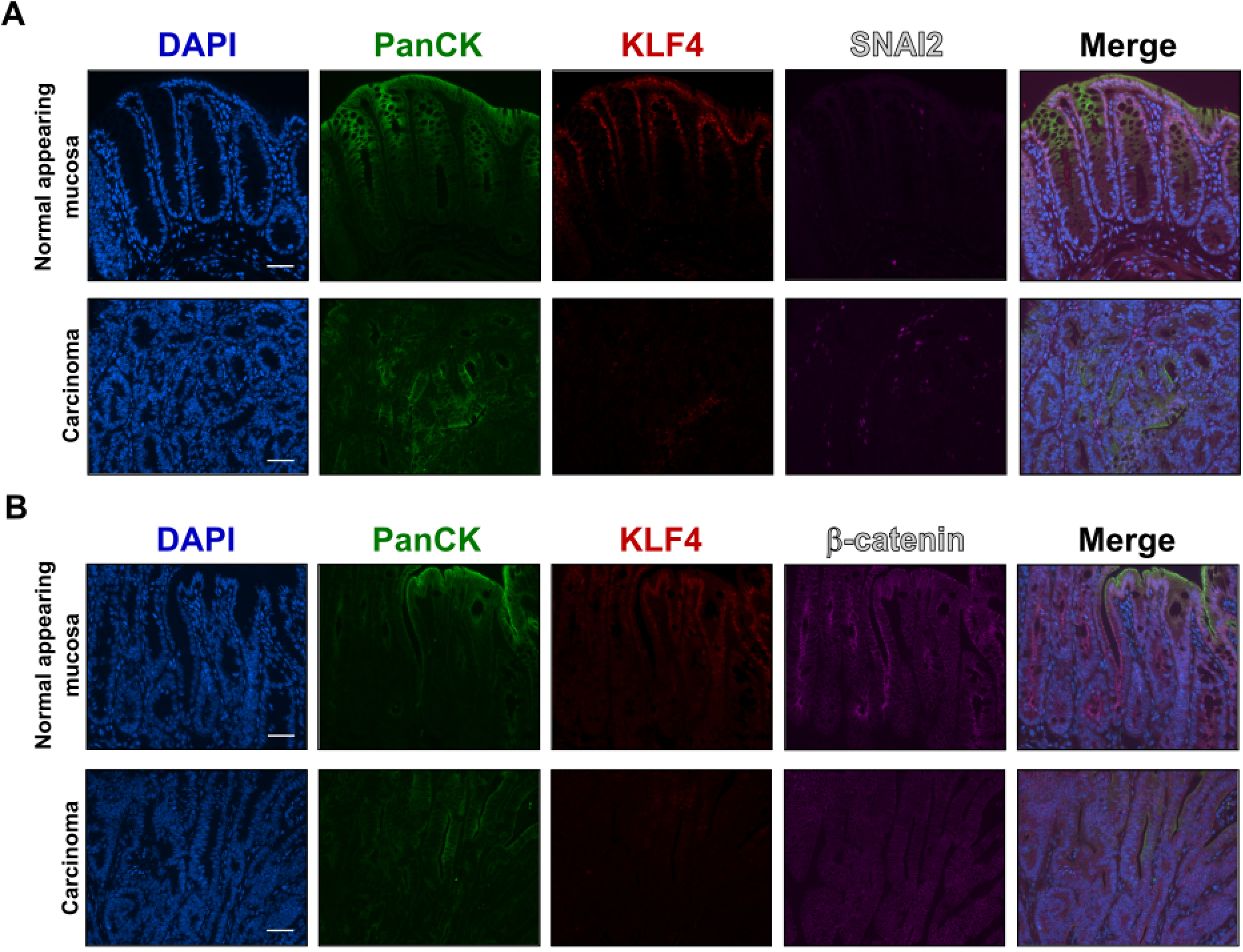
Immunofluorescence staining for PanCK, KLF4, SNAI2 and, β-catenin in mouse colonic tissues after AOM/DSS treatment. (**A**) Representative images of PanCK, KLF4, SNAI2 in normal adjacent mucosa (top panel) and tumor sections (bottom panel). (**B**) Representative images of PanCK, KLF4, and β-catenin in normal adjacent mucosa (top panel) and tumor sections (bottom panel). Scale bar, 50 μm.

**Figure 5.**
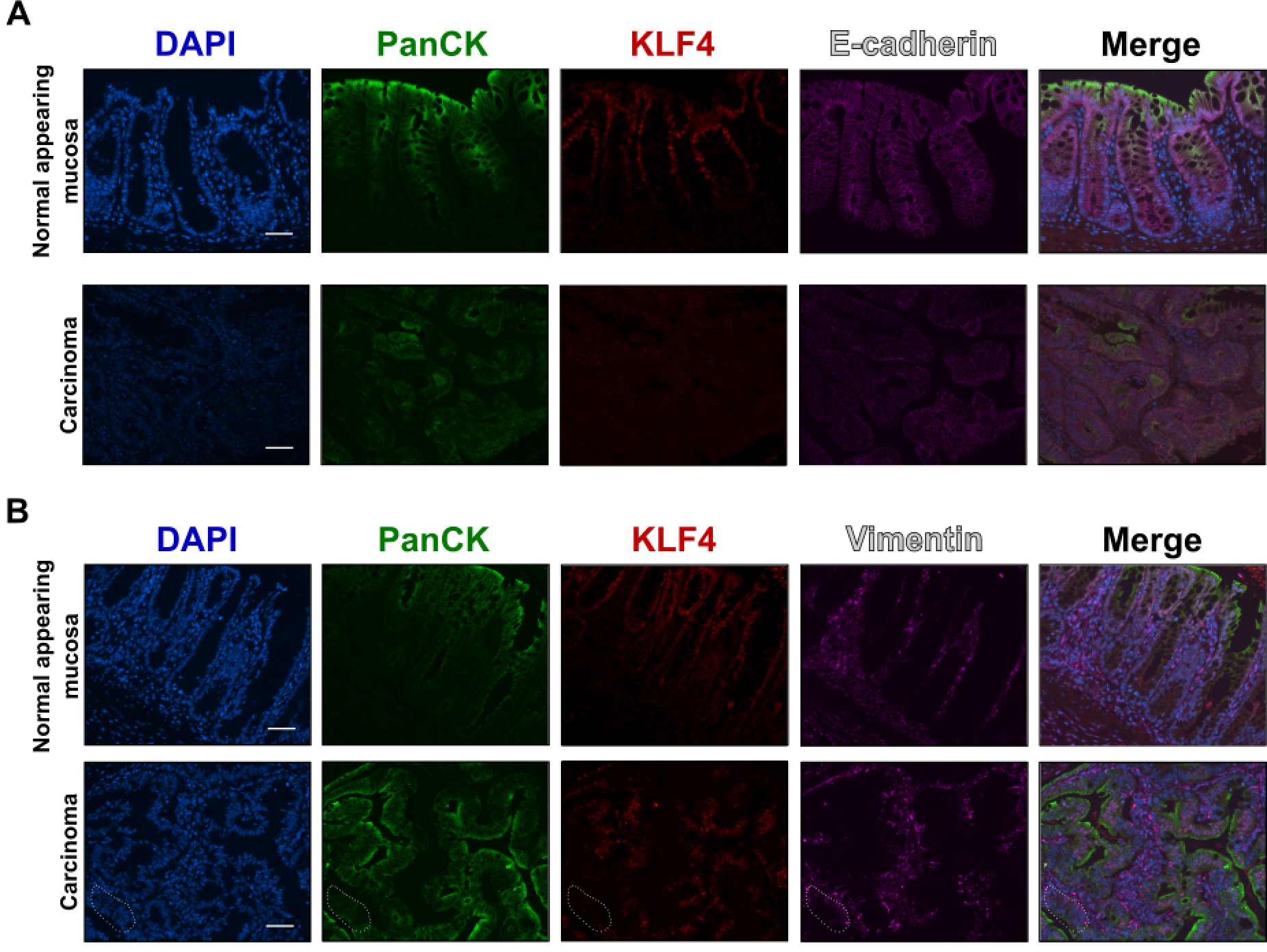
Immunofluorescence staining for PanCK, KLF4, E-cadherin, and vimentin in mouse colonic tissues after AOM/DSS treatment. (**A**) Representative images of PanCK, KLF4, and E-cadherin in normal adjacent mucosa (top panel) and tumor sections (bottom panel). (**B**) Representative images of PanCK, KLF4, and vimentin in normal adjacent mucosa (top panel) and tumor sections (bottom panel). Dotted area shows transition from epithelial (PanCK staining) to mesenchymal characteristic (vimentin staining). Scale bar, 50 μm.

## Discussion

In this paper, we investigated the correlation in expression between KLF4 and EMT markers in tissues obtained from patients with colorectal cancer (CRC) and from a mouse model of colitis-associated cancer. The role of KLF4 in the regulation of EMT in cancer is debatable. The results from studies of epidermal cancer, hepatocellular carcinoma, breast cancer, pancreatic cancer, and prostate cancer with data predominantly originating from *in vitro* experiments show that KLF4 negatively regulates EMT (28–33). On the other hand, KLF4’s ability to regulate the stemness of cancer cells has been shown as an important factor in stimulating EMT in pancreatic, ovarian, endometrial, nasopharyngeal, prostate, and non-small lung cancers (34–40). Here, we demonstrate that KLF4 expression is positively correlated with epithelial markers of EMT in normal mucosa and negatively with mesenchymal markers in CRC. These results are in agreement with previous observations that KLF4 is a suppressor of EMT (28–31, 41). This could be accredited to the role of KLF4 in the regulating differentiation along the crypt-luminal axis in the intestinal epithelium (10). Furthermore, studies using conditional ablation of *Klf4* from the intestinal epithelium showed a deficiency in goblet cell differentiation. Thereby demonstrating that KLF4 plays a role in maintaining intestinal epithelial morphology and homeostasis (9, 12). Additionally, KLF4 has been shown to regulate apical-basolateral polarity in intestinal epithelial cells and to enhance their polarity by transcriptional regulation (13). Thus, loss of KLF4 expression during development and progression of CRC may lead to loss of cell polarity with progression toward EMT. Furthermore, it has been previously demonstrated that E-cadherin (*Cdh1*), N-cadherin (*Cdh2*), vimentin (*Vim*), and β-catenin (*Ctnnb1*) genes are direct transcriptional targets of KLF4 (30). Yori and colleagues demonstrated that KLF4 is necessary for the maintenance of E-cadherin expression, repression of SNAI1 and prevention of EMT in mammary epithelial cells (29, 41). These results are consistent with our observations. We demonstrated a positive correlation between KLF4 and E-cadherin and a negative one between KLF4 and N-cadherin and vimentin in human and mouse tissues. However we were not able to show increased nuclear levels of β-catenin in mouse tissues after AOM/DSS treatment (18). This could be due to technical differences, as previously we used immunohistochemical staining and the current analysis was based on immunofluorescence studies. Also, we showed that KLF4 expression is negatively correlated to the expression of TWIST, SNAI2, and claudin-1. Taken together, these results suggest that KLF4 may regulate EMT at several different stages of CRC progression, including suppression of expression of transcription factors (TWIST and SNAI2), proliferation markers (β-catenin) and regulation of cell polarity (E-cadherin, N-cadherin, and claudin-1). In conclusion, we showed for the first time a negative association between KLF4 and mesenchymal EMT markers in both human and mouse CRC tissues. Future studies are necessary to identify the mechanism by which KLF4 regulates EMT progression in CRC.

## Acknowledgments

We thank the Department of Pathology at Stony Brook University for technical assistance in histopathological analysis. K.C.G, J.Z.H, and A.M.G conducted experiments and analyzed data; J.L.W provided tissues samples; K.R.S assisted with histopathological analysis; V.W.Y, A.M.G, and A.B.B conceived the main idea and revised the paper. All authors approved final version of the paper. This work was supported by grants from the National Institutes of Health awarded to V.W.Y. (CA084197).

